# Real-time sharing of drug screening data on the NCATS OpenData Portal accelerates translational research

**DOI:** 10.1101/2020.06.04.135046

**Authors:** Kyle R. Brimacombe, Peter C. Scully, Tongan Zhao, Richard T. Eastman, Xin Hu, Ke Wang, Mark Backus, Bolormaa Baljinnyam, Catherine Z. Chen, Lu Chen, Yichen Cheng, Danielle P. Davis, Kevin Duerr, Tara Eicher, Aaron Friedman, Ying Fu, Kirill Gorshkov, Hui Guo, Quinlin M. Hanson, Nathan Hotaling, Zina Itkin, Stephen C. Kales, Carleen Klumpp-Thomas, Emily M. Lee, Sam Michael, Tim Mierzwa, Brian Nezhad, Andrew Patt, Brittany J. Poelaert, Manisha Pradhan, Alex Renn, Paul Shinn, Jonathan H. Shrimp, Therese N. Tripler, Amit Viraktamath, Kanny K. Wan, Kelli M. Wilson, Miao Xu, Alexey V. Zakharov, Wei Zhu, Wei Zheng, Anton Simeonov, Ewy A. Mathé, Marc Ferrer, Donald C. Lo, Min Shen, Matthew D. Hall

## Abstract

The National Center for Advancing Translational Sciences (NCATS) has developed an online open science platform – named the NCATS OpenData Portal (ODP) – for quickly and freely sharing complete NCATS translational datasets via an open-access, user-friendly interface. This paper describes the establishment of the ODP, initially deployed during the COVID-19 crisis, and provides a detailed analysis of COVID-19 drug repurposing screening datasets that served as the first large-scale use case for the platform. Over 10,000 compounds were tested across 17 quantitative high-throughput assays, covering a wide spectrum of the SARS-CoV-2 life cycle. In total, over 87,000 concentration-response curves and 426,000 data points were made publicly available on ODP in near real-time, enabling immediate access to complete datasets. The resource is flexible in accommodating various types and structures of data, and it has already expanded since its launch to host additional datasets for COVID-19, other viruses of pandemic potential, and beyond. The OpenData Portal has been designed as a scalable platform for real-time data sharing across drug discovery campaigns, regardless of disease area, with the overarching goal of accelerating discovery at NCATS, the NIH, and the greater scientific community.

## Introduction

On March 11, 2020, the World Health Organization declared the outbreak of atypical pneumonia coronavirus disease (COVID-19) - caused by the novel betacoronavirus SARS-CoV-2 - to be a pandemic^1^. Spurred by the significant public health and economic consequences of the virus, biomedical scientists around the world at academic and government laboratories, biotechnology companies, and pharmaceutical corporations rapidly mobilized to understand the disease and develop protective vaccines and therapeutic interventions to mitigate its impact. Clinical trials were quickly undertaken for many drug repurposing candidates, notably remdesivir^2^, which received Emergency Use Authorization (EUA) from the Federal Drug Agency (FDA) in the United States on May 1, 2020^3^, and dexamethasone, which was adopted into practice in the United Kingdom on June 16, 2020^4^. Drugs were initially prioritized for clinical trials based on 1) anecdotal clinical observations, 2) prior history as antivirals, and 3) a contemporary yet evolving understanding of SARS-CoV-2 biology. In parallel, repurposing screens were implemented across the research community in search of unrealized SARS-CoV-2 activity possessed by approved and experimental agents^5–13^.

Recognizing the lack of public, comprehensive SARS-CoV-2 screening datasets (including inactive compound data), and the need for quick and open sharing, the National Center for Advancing Translational Sciences (NCATS) began developing the OpenData Portal (ODP) in the Spring of 2020. This online resource was created to facilitate the rapid sharing of complete datasets from NCATS’s own large panel of target-based SARS-CoV-2 screening campaigns. Although the ODP was built in response to the COVID-19 pandemic, it was designed from the start to provide a scalable, established platform that would enable data-sharing for future screens, regardless of disease. The OpenData Portal is also flexible in accommodating various types and structures of data, and it has expanded to host additional datasets beyond screening data. These new data sets include curated metadata on *in vivo* COVID-19 studies, a catalog of animal models established for SARS-CoV-2 research, and a vast curated database of therapeutic activity reported against known and emerging SARS-CoV-2 variants, among others. The following analysis and description are focused on the original objective of the OpenData Portal, which was rapid sharing of complete drug repurposing screening datasets for SARS-CoV-2 to accelerate translational research efforts.

## Results

### OpenData Portal Establishment and Goals

The OpenData Portal dashboard was first shared publicly on May 25, 2020, with the goal of accelerating the development of COVID-19 therapeutics. NCATS was engaged in a coordinated effort to create a panel of novel assays to probe the SARS-CoV-2 viral replication cycle and generate large quantities of activity data as quickly as possible. The specific objective of the OpenData Portal was to inform COVID-19 research hypotheses and support further mechanistic studies of compounds capable of inhibiting viral infectivity^14^, by providing the global research community with access to these validated methodologies and actionable data in close to real-time. To be successful, the OpenData Portal would need to share datasets that were both more complete - including counterscreen and negative results - and more rapidly available than via traditional publication processes.

Meeting these initial objectives required the following: 1) a set of validated high-throughput assays against SARS-CoV-2 targets, 2) libraries of compounds with potential activity against those targets, 3) the resources and infrastructure to conduct quantitative high-throughput screening campaigns using those libraries, and 4) an open-access, user-friendly interface to rapidly disseminate the screening results.

### SARS-CoV-2 Assay Development

The SARS-CoV-2 assays developed at NCATS and made available via the OpenData Portal were designed to probe each phase of the viral replication cycle, covering a wide array of both viral and human (host) targets (**Figure 1a**). In total, seventeen validated assay protocols were shared on the OpenData Portal (**Table 1**). They are grouped into the following four target classes based on different mechanisms of experimental design (**Figure 1b**): 1) **live virus infectivity** assays (2 out of 17 total assays) measure the ability of live SARS-CoV-2 virus to infect and replicate within human host cells *in vitro*; 2) **pseudovirus infectivity** assays (6/17) use simplified, recombinant viral particles to characterize initial binding and entry of virus into human host cells *in vitro* at reduced biocontainment levels; 3) **viral entry** assays (5/17) measure the activity of targets related to virion binding and entry (e.g. viral spike protein, human ACE2, human TMPRSS2); and 4) **viral replication** assays (2/17) measure the activity of viral enzymes necessary for genome processing and replication (e.g., 3CL protease and RNA-dependent RNA-polymerase (RdRp)). Assays are also grouped according to type (**Figure 1c**): **cell viability** assays (7/17) quantifying relative numbers of live host cells to determine anti-proliferative effects of treatment with virus and/or compound (e.g. SARS-CoV-2 cytopathic effect (CPE)); cell-based **reporter gene** assays (3/17) utilizing recombinant viral constructs to measure expression of reporter proteins like luciferase (e.g. SARS-CoV-2 pseudotyped particle entry); protein-based **proximity**/binding assays (2/17) (e.g. spike-ACE2 protein-protein interaction (AlphaLISA)); a **phenotypic** high-content imaging-based assay (1/17) measuring cellular localization and internalization of human ACE2 receptor in host cells in the presence of spike-receptor binding domain protein tagged to quantum dots (spike-ACE2 protein-protein interaction (Quantum Dot)); and **biochemical** assays (4/17) measuring enzymatic activity of SARS-CoV-2 or human targets (e.g. SARS-CoV-2 3CL protease and human TMPRSS2 enzyme). Most assays utilized a luminescent reporter/readout (10/17), followed by fluorescence (5/17) and AlphaLISA (2/17) (**Figure 1d**).

**Figure 1.**
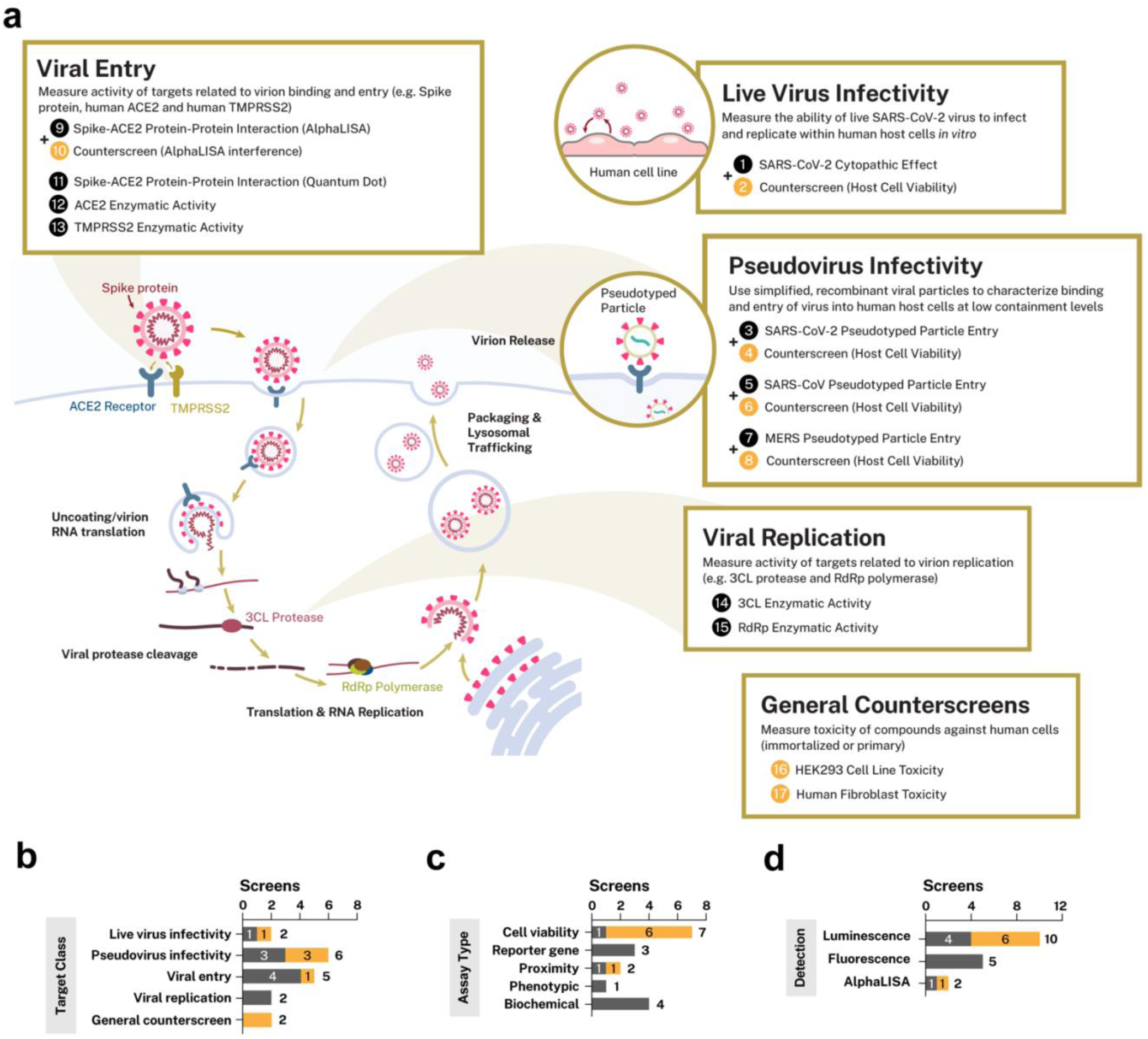
NCATS viral high-throughput screening assays shared via the OpenData Portal, shown in the context of the SARS-CoV-2 viral replication cycle. **a)** SARS-CoV-2 screening assays with available data are shown organized by target class. Assays are numbered to match Table 1; primary assays are labeled in black and paired counterscreens and general counterscreens are labeled in orange. Where applicable, paired counterscreen assays (i.e. those run to identify artifacts in specific screens) are shown next to their matched primary assay. Tan trails indicate the portion of the SARS-CoV-2 replication cycle that each target class is designed to interrogate, where applicable. Bar charts illustrating the distribution of NCATS SARS-CoV-2-related screening campaigns are shown by **b)** target class, **c)** assay type and **d)** detection technology. Within each category, bars are broken out into primary screening assays (grey) and counterscreens (orange).

**Table 1.** SARS-CoV-2-related screening assays in the NCATS OpenData Portal. Summary of the validated SARS-CoV-2 assays that were shared on the OpenData Portal, including the small molecule libraries that each assay was screened against.

| Assay Name | Target Class | Assay Type | Detection Type | Assay Description | NPC | N<br>P<br>A<br>C<br>T | M<br>I<br>P<br>e | A<br>n<br>t<br>i-<br>i<br>n<br>f<br>e<br>c<br>t<br>i<br>v<br>e |
| --- | --- | --- | --- | --- | --- | --- | --- | --- |
| SARS-CoV-2 Cytopathic Effect | Live virus infectivity | Cell viability | Luminescence | Measures ability of compounds to mitigate viral-induced cytopathic effect (CPE) against host cells | X | X | X | X |
| SARS-CoV-2 Cytopathic Effect (Tox Counter) | Live virus infectivity | Cell viability | Luminescence | Detects compound toxicity against host cells in absence of virus | X | X | X | X |
| SARS-CoV-2 Pseudotyped Particle Entry | Pseudovirus infectivity | Reporter gene | Luminescence | Measures inhibition of recombinant SARS-CoV-2 viral particle cell entry in cells using reporter gene expression | X |  | X |  |
| SARS-CoV-2 Pseudotyped Particle Entry (Tox Counter) | Pseudovirus infectivity | Cell viability | Luminescence | Detects compound toxicity against host cells in absence of pseudovirus | X |  | X |  |
| SARS-CoV Pseudotyped Particle Entry | Pseudovirus infectivity | Reporter gene | Luminescence | Measures inhibition of recombinant SARS-CoV viral particle cell entry in cells using reporter gene expression | X |  |  |  |
| SARS-CoV Pseudotyped Particle Entry (Tox Counter) | Pseudovirus infectivity | Cell viability | Luminescence | Detects compound toxicity against host cells in absence of pseudovirus | X |  |  |  |
| MERS Pseudotyped Particle Entry | Pseudovirus infectivity | Reporter gene | Luminescence | Measures inhibition of recombinant MERS viral particle cell entry in cells using reporter gene expression | X |  |  |  |
| MERS Pseudotyped Particle Entry (Tox Counter) | Pseudovirus infectivity | Cell viability | Luminescence | Detects compound toxicity against host cells in absence of pseudovirus | X |  |  |  |
| Spike-ACE2 Protein-Protein Interaction (AlphaLISA) | Viral entry | Proximity | AlphaLISA | Measures disruption of binding between SARS-CoV-2 spike protein and ACE2 host receptor | X |  |  | X |
| Spike-ACE2 Protein-Protein Interaction (AlphaLISA Counter) | Viral entry | Proximity | AlphaLISA | Measures off-target inhibition of AlphaLISA detection technology | X |  |  | X |
| Spike-ACE2 Protein-Protein Interaction (Quantum Dot) | Viral entry | Phenotypic | Fluorescence | Measures internalization/localization of human ACE2 receptor in host cells in the presence of quantum dot-tagged spike-RBD protein | X* | X* | X* | X* |
| ACE2 Enzymatic Activity | Viral entry | Biochemical | Fluorescence | Measures inhibition of recombinant human ACE2 catalytic activity | X |  |  | X |
| TMPRSS2 Enzymatic Activity | Viral entry | Biochemical | Fluorescence | Measures inhibition of recombinant human TMPRSS2 protease activity | X |  | X |  |
| 3CL Enzymatic Activity | Viral replication | Biochemical | Fluorescence | Measures inhibition of recombinant SARS-CoV-2 3CL protease activity | X | X | X | X |
| RdRp Enzymatic Activity | Viral replication | Biochemical | Fluorescence | Measures inhibition of recombinant SARS-CoV-2 RdRp activity | X |  |  |  |
| HEK293 Cell Line Toxicity | General counterscreen | Cell viability | Luminescence | Detects compound toxicity against HEK293 cells | X | X |  |  |
| Human Fibroblast Toxicity | General counterscreen | Cell viability | Luminescence | Detects compound toxicity against human fibroblasts | X |  | X |  |

\*Tested at a single concentration

The OpenData Portal also includes data from counterscreen assays, which help to identify and rule out potential false positives caused by artifacts such as assay interference or cytotoxic effects. Five of the primary assays are paired with a counterscreen assay (**Figure 1**; **Table 1**), which measures either off-target toxicity against host cells lines or potential interference with assay detection technologies (e.g. AlphaLISA), to aid interpretation of activity seen in the corresponding primary assays. There are also two **general counterscreens** included that report the effects of screening compounds on human cell growth and viability in general, against an immortalized cell line and primary fibroblasts, respectively. These general counterscreens were intended to give a broader sense of bioactivity of these compounds in human cells and provide additional context for activity signatures seen across cell-based assays.

Importantly, all assay documentation including assay overviews, methodology and detailed assay protocols were instantly made available and readily retrievable on the ODP website alongside the screening results to facilitate adoption and ensure that other laboratories can replicate these assays (https://opendata.ncats.nih.gov/covid19/assays).

### Compound Libraries

To maximize the potential for drug repurposing and accelerate therapeutic development, screening efforts prioritized four annotated small molecule libraries (**Table 2**). These libraries included 1) the NCATS Pharmaceutical Collection (NPC), a library containing most compounds approved for use by the Food and Drug Administration, late stage clinical candidates, and drugs approved by regulatory agencies in other countries^15,16^; 2) the NCATS Pharmacologically Active Chemical Toolbox (NPACT) library and 3) Mechanism Interrogation Plates (MIPe) library, two annotated/bioactive libraries of diverse small molecules with known mechanisms-of-action, including small molecule probes and experimental therapeutics designed to modulate a wide range of targets^17,18^; and 4) a library of 732 anti-infective compounds with potential anti-coronavirus activity that have been reported in the literature, which was assembled at NCATS specifically for COVID-19 screening efforts and is published here for the first time (**Supplemental Table 1**). Collectively, these libraries contain 10,803 compounds; there is a small degree of overlap among these libraries (13.8%), leaving 9,312 structurally unique compounds. Of these, 1,820 (20%) are approved for use in humans, and an additional 989 (11%) had entered human clinical evaluation (Phase 1-3 trials) at the time of testing. Importantly, 5,913 (63%) of these compounds have at least one annotated target, providing valuable insight into potential mechanisms-of-action if any of those compounds were found to perturb the SARS-CoV-2 life cycle.

**Table 2.** Small molecule libraries screened and included in the NCATS OpenData Portal. Description of the small molecule libraries that were screened and used to generate the activity data shared on the OpenData Portal.

| Small Molecule Library | Library Type | Size | Description | Publication(s) |
| --- | --- | --- | --- | --- |
| NCATS Pharmaceutical Collection (NPC) | Approved drugs | 2,678 compounds | The NCATS Pharmaceutical Collection (NPC) is a library containing most compounds that have been approved for use by the Food and Drug Administration, late-stage clinical candidates, and a number of approved molecules from related agencies in foreign countries. | 15,16 |
| NCATS Pharmacologically Active Chemical Toolbox (NPACT) | Annotated / bioactive compounds | 4,916 compounds | The NCATS Pharmacologically Active Chemical Toolbox (NPACT) is a library of annotated compounds that inform on novel phenotypes, biological pathways and cellular processes. There are more than 7,000 mechanisms and phenotypes identified in the literature and worldwide patents that cover biological interactions within mammalian, microbial, plant and other model systems. | 70 |
| Mechanism Interrogation Plate (MIpe) | Annotated / bioactive compounds | 2,477 compounds | Originally designed as an oncology-focused library, NCATS's Mechanism Interrogation Plate (MIpe) is a library of pharmacologically defined small molecules with relevance to cancer, neurology, infectious diseases and stem cell biology. | 70,71 |
| Anti-infective Library | Anti-infective | 732 compounds | This library was curated by NCATS early in the pandemic, and contains compounds reported in the literature with anti-infective activity or identified as potential repurposing candidates for coronavirus infection. | Supplemental Table 1 |

### Summary of SARS-CoV-2 Screening Efforts

Seventeen screening campaigns, against either SARS-CoV-2 targets or associated counterscreens, have been completed to date (**Table 1**), and the full datasets, comprising nearly half a million data points, have been made publicly available via the OpenData Portal (**Supplemental Table 2**). The use of quantitative high-throughput screening (qHTS) allowed concentration-response curves to be fit to nearly all primary screening data^19^. Each SARS-CoV-2 screen generated between 2,600-11,000 compound concentration-response curves each, with over 85,000 curves generated and shared in total.

Reagent supply was a limiting factor for many assays early in the pandemic, so drug-repurposing screening was generally prioritized to maximize the potential for new therapeutics. To this end, nearly all SARS-CoV-2 assays were screened against the NPC approved drugs collection at a minimum of 4 concentrations (16/17), and most of those (11/16) were tested against at least one additional library (**Table 1**). Due to its high resource cost, the high-content spike-ACE2 quantum dot assay was run at just a single concentration against the NPC, NPACT, MIPE and anti-infective libraries; identified hits (representing 120 compounds) were plated in concentration-response and retested in triplicate.

Details of the data analysis, curve fitting, and classification methodology are provided in the Methods section below. Briefly, raw screening results were normalized to assay-relevant controls, and quality control metrics were calculated for each normalized plate. Three- or four-parameter curves were fit to all concentration-response series, and curves were classified according to the magnitude of response (efficacy), quality of the curve fit to the data (r^2^), and the number of asymptotes to the calculated curve. Compounds assigned a curve class of 1.1, 1.2, 2.1 or 5 were categorized as high confidence actives, those assigned curve classes of 1.3, 1.4, 2.2, 2.3, 2.4 or 3 were categorized as low confidence actives, and the remainder were categorized as inactives.

Total hit rates for the SARS-CoV-2 screens (i.e. percentages of compounds in either high or low confidence active category) ranged from 0.4% to 44% (**Figure 2a**), and across all assays, 5.3% of compounds (4,566) demonstrated high confidence activity, 13.3% showed low confidence activity (11,346) and 81.4% (69,973) were found to be inactive (**Figure 2b**). Biochemical assays yielded lower hit rates on average (3%), while screens utilizing reporter gene expression or proximity measurements (e.g. AlphaLISA) as readouts gave the highest average hit rates (35% and 31%, respectively). The high-content spike-ACE2 quantum dot screen was excluded from the hit rate analysis, since the primary screen was run at a single concentration; however, concentration-response data was generated for a subset of cherrypicked compounds showing activity (specifically, calculated relative spot intensity data) and loaded onto the OpenData Portal.

**Figure 2.**
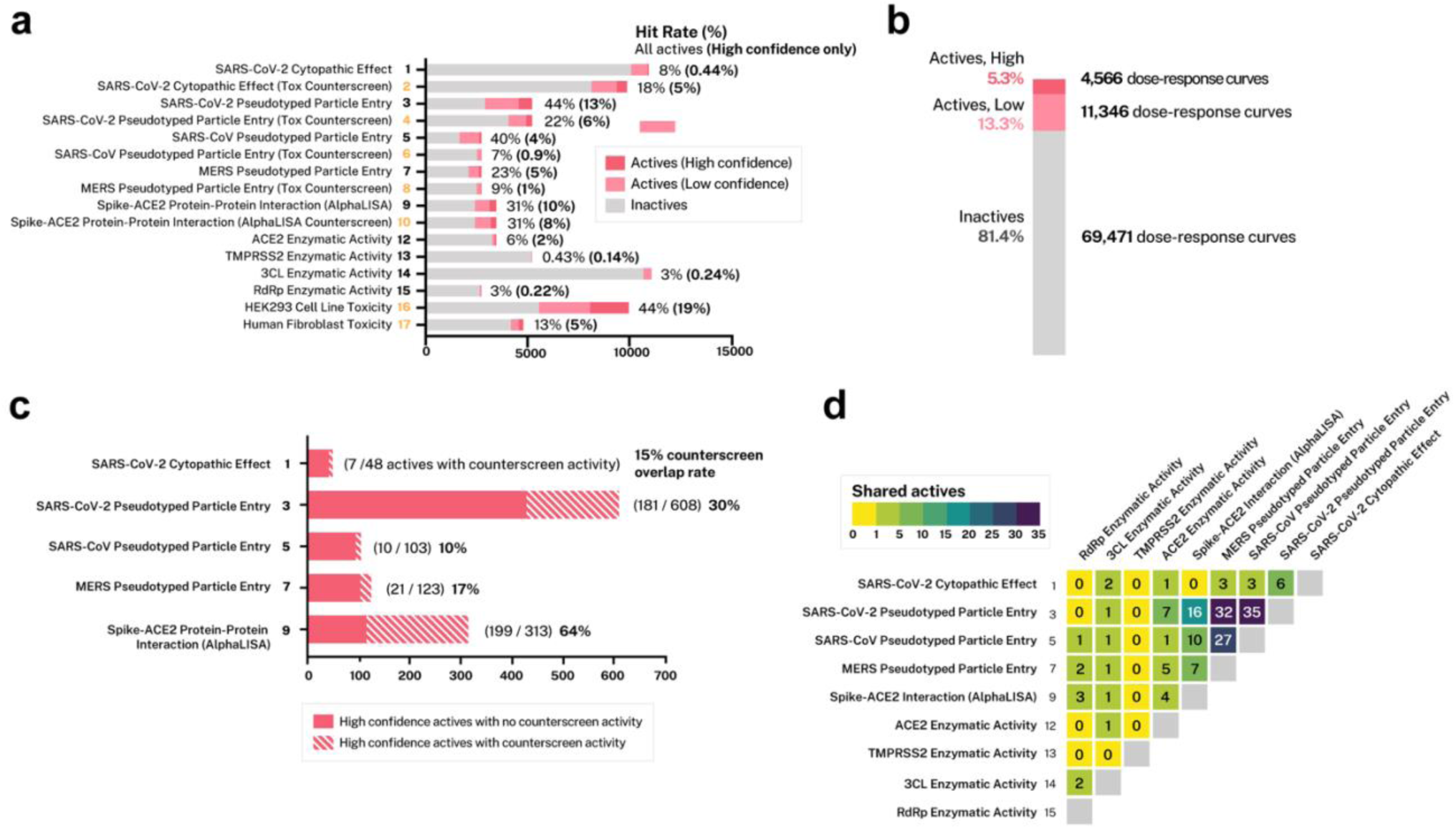
Summary of SARS-CoV-2 high-throughput screening campaigns and data. **a**) Activity breakdown for individual SARS-CoV-2 high-throughput screens, illustrating the proportion of compounds that were scored as inactives (grey), low confidence actives (pink) and high confidence actives (dark pink). **b**) Cumulative activity breakdown for all SARS-CoV-2 high-throughput screens, indicating proportion of inactives (grey), low confidence actives (pink) and high confidence actives (dark pink). **c**) Bar chart illustrating proportions of high confidence actives from primary assays exhibiting high confidence activity in a paired counterscreen assay, by screen (where applicable). **d**) Heatmap depicting the number of active compounds shared between each pair of primary SARS-CoV-2 screens; for this portion of the analysis, active compounds are those that 1) scored as high confidence actives in a primary screen and 2) did not demonstrate high confidence activity in the corresponding counterscreen (where applicable).

Of the SARS-CoV-2 assays screened, five utilized paired counterscreen assays to identify potential artifacts or compounds with activity that could confound interpretation of the primary screen. Specifically, four of the five paired counterscreen assays measured viability of the host cell line in the absence of virus or pseudovirus; these screens were designed to highlight compounds capable of effecting toxicity or growth inhibition on the host line (which can confound data normalization and interpretation), rather than on the virus or virus-surrogate directly. The fifth paired counterscreen was a TruHits assay (Revvity, Waltham, MA) specifically designed to identify compounds capable of interfering with the AlphaLISA proximity detection technology, mimicking inhibition in the absence of the intended spike and ACE2 protein targets. The SARS-CoV and MERS pseudovirus assays and the live virus SARS-CoV-2 CPE assay showed the lowest hit rates in their paired counterscreen assays, with 10% to 17% of primary screen hits showing high confidence activity (**Figure 2c**). The SARS-CoV-2 pseudovirus assay demonstrated a 30% counterscreen hit rate (181 out of 608 total hits), while 64% of primary hits in the spike-ACE2 AlphaLISA assay were active in the TruHits counterscreen (**Figure 2c**).

Nearly all of NCATS’s SARS-CoV-2 assays were screened against the NPC approved drugs library, allowing for an assessment of overlap in hits between the individual assays (**Figure 2d**). After removing compounds with activity in a paired counterscreen assay, the greatest overlap in screening hits was seen between the three pseudovirus screens, which shared 27-35 hits in common. The spike-ACE2 AlphaLISA assay also showed overlap with the SARS-CoV-2 pseudovirus assay (16 hits in common), and modest overlap with the other two pseudovirus screens as well (7-10 hits in common). The enzymatic screens - notably the 3CLpro, RdRp and TMPRSS2 assays - ranged from 0 to 3 hits in common.

Due to the small degree of overlap among the compound libraries noted above, some compounds were tested in the same SARS-CoV-2 assays multiple times, generating replicate data. As a further measure of assay performance and reproducibility, the activity of these replicate compounds was examined. For the purposes of this analysis, “confirmed” replicates were defined as compounds that demonstrated activity of the same category (i.e. high confidence active, low confidence active, inactive) in all instances when tested against the same screening assay. Observed internal confirmation rates were high among all assays, ranging from 70.6% to 99.68% of replicates confirmed (**Table 3**).

**Table 3.**
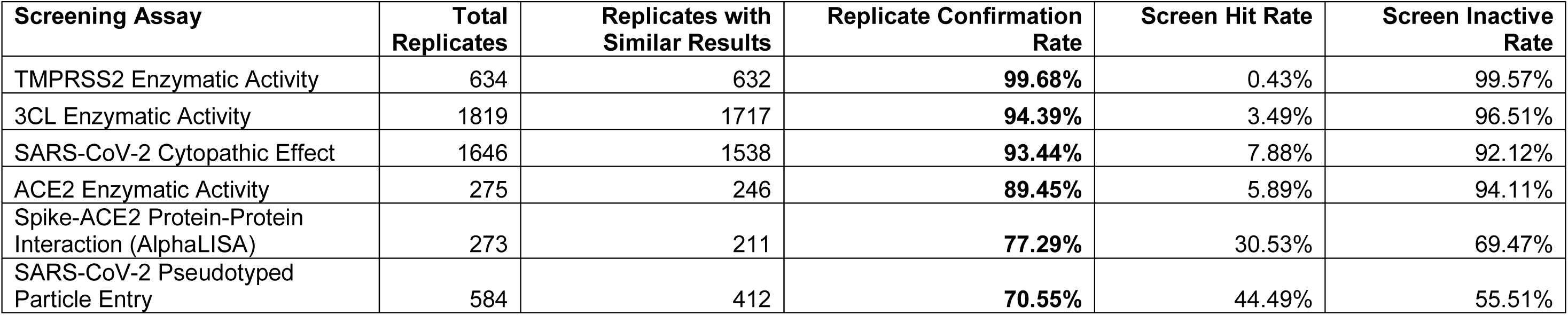
Confirmation rate of duplicated compounds in NCATS screens. Summaries of the number of duplicate compounds screened and the percentage that confirmed are shown for each screening assay, along with the overall screen hit rate.

### Speed of Screening Data-sharing

High-throughput screening for the first set of SARS-CoV-2 target assays began in April 2020, with most screening efforts taking place between May 2020 and March 2021. Consistent with the mission of the OpenData Portal, the majority of the SARS-CoV-2 complete screening datasets were shared on the site in advance of preprint or peer-reviewed publication release, with 59% (10/17) shared before publication and another 29% (5/17) shared within 2 weeks of publication (**Figure 3a**). Most of this data was released within the first year of the COVID-19 pandemic; five screens were shared at the launch of the OpenData Portal in May 2020, 8 screens were shared during the remainder of 2020, and 4 were posted in 2021 and 2022 (**Figure 3b**).

**Figure 3.**
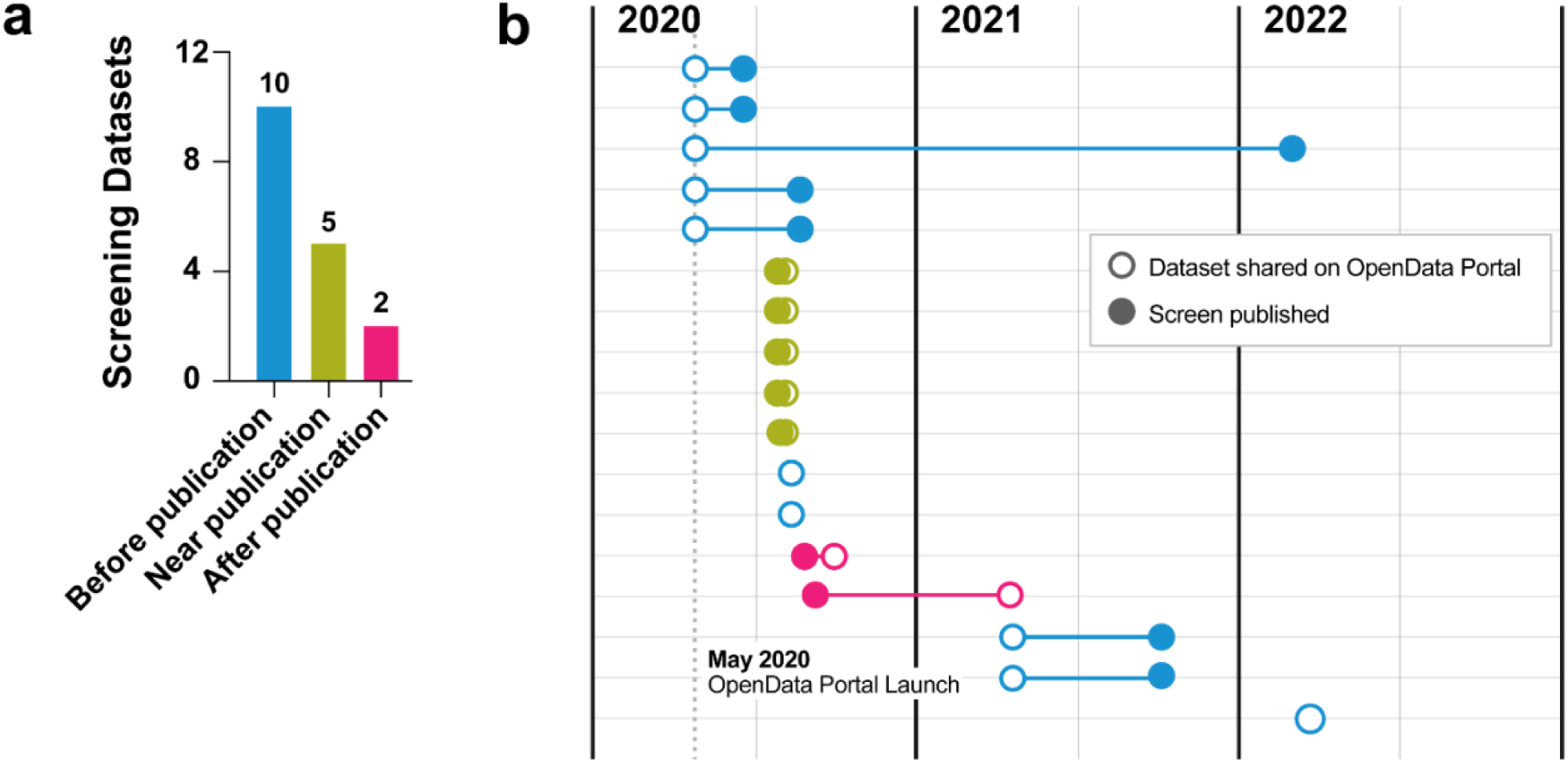
Rapid data sharing of OpenData Portal SARS-CoV-2 high-throughput screening campaign data. **a)** Bar chart illustrating the number of NCATS SARS-CoV-2 screening datasets that were publicly shared on OpenData Portal before (blue), near (green) or after (magenta) publication of their respective screening campaigns. **b)** Dumbbell chart showing a timeline of release dates for the 17 screening datasets on OpenData Portal. Open circles indicate dates of public data releases on OpenData Portal, while filled circles indicate publication dates for the associated assay screening efforts. Series colors indicate whether the assay data was shared before (blue), near (green) or after (magenta) publication.

### Design of the SARS-CoV-2 Screening Data Browser

A key goal of the OpenData Portal website was to allow users to explore NCATS’s SARS-CoV-2 screening data in an intuitive way and help guide the exploration of new therapeutic hypotheses. The site was built to emphasize searchability by a general science audience, featuring a user-friendly heatmap visualization^20^ to allow direct and convenient comparison of compound activity across the panel of all assays tested, including orthogonal and counterscreen assays (**Figure 4a**). Within the heatmap, green squares denote activity in a primary screen, while pink squares indicate activity in a relevant counterscreen, alerting users to cases where primary activity should be interpreted with caution due to potential off-target activity. Darker colors indicate compounds that are more potent or efficacious (high confidence actives) and lighter colors indicate less potent or efficacious compounds (low confidence actives).

**Figure 4.**
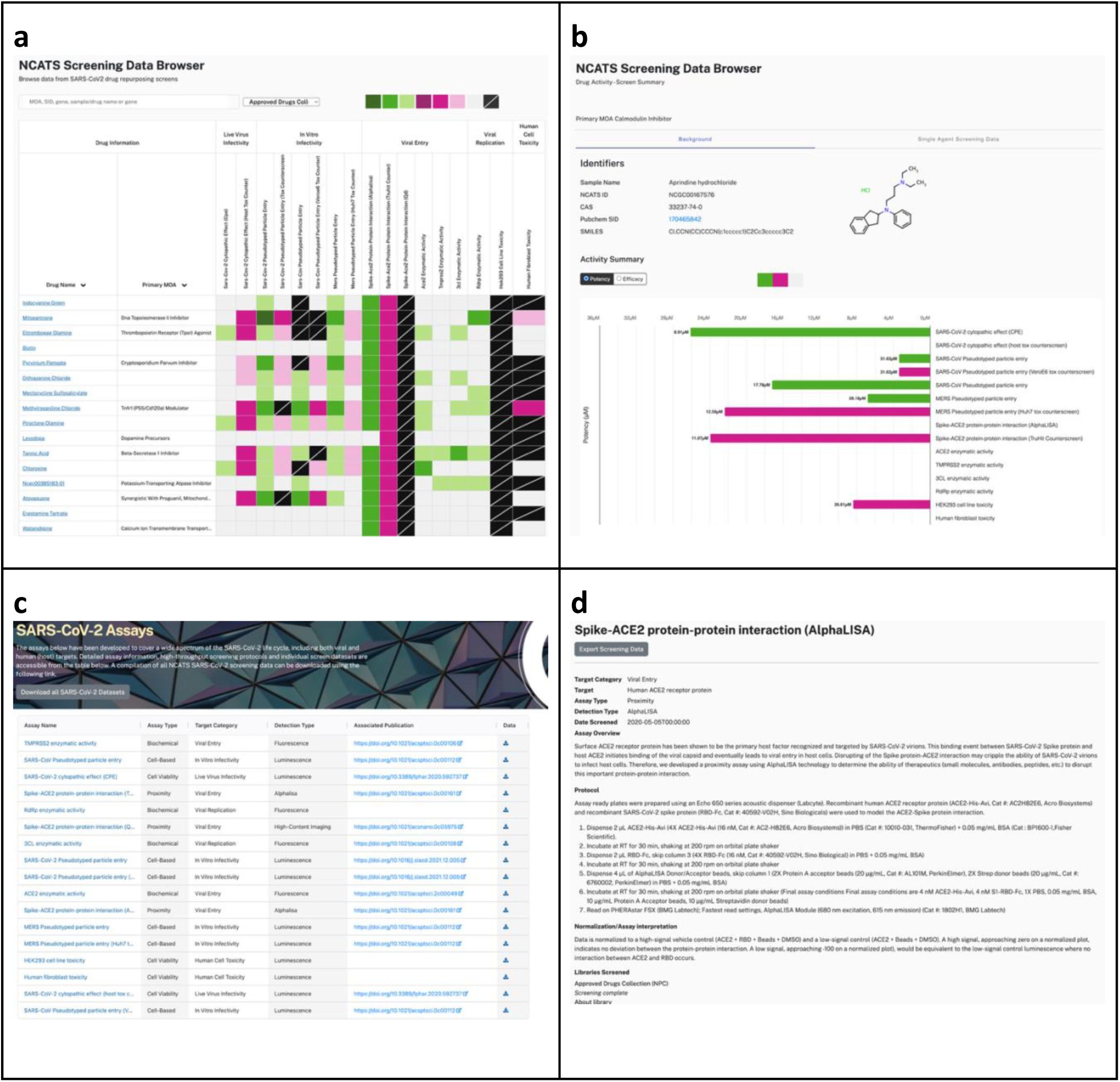
OpenData Portal enables exploration of high-throughput screening data. **a**) Screening Data Browser page, with heatmap visualization of activity of compounds (rows) across various assays (columns), for direct comparison. Green squares denote activity in a primary screen, while pink squares indicate activity in a relevant counterscreen; darker colors indicate compounds that are more potent or efficacious (high confidence actives) and lighter colors indicate less potent or efficacious compounds (low confidence actives). Grey squares denote inactive compounds, and black squares indicate that a compound was not included in a screen. The browser has both sort and search functionalities to facilitate interrogation of data. **b**) Individual compound activity pages, which summarize activity (both potency and efficacy) for single compounds across all SARS-CoV-2 assays screened, along with graphs displaying concentration-response data and curves. Compound structure and metadata are also provided. **c**) Assay Overview Page organizes access to all NCATS SARS-CoV-2 assays, including methodology and detection strategy, as well as links to download the complete screening dataset for each assay **d**) Individual assay pages, which provide detailed assay protocols to allow independent evaluation and confirmation of method and compound activities, as well as assay performance across different screening libraries.

A search box allows users to filter the displayed compounds based on a variety of different data classes, including drug name, compound ID, primary mechanism-of-action (where known), and target gene ID. Users can also re-sort the heatmap to rank compounds by their activity within a specific assay of interest. By clicking on individual compounds, users can access a summary-level view with more detailed information, including graphs with concentration-response data and curve fits for all relevant screening assays for that compound (**Figure 4b**).

The website also contains an Assay Overview Page where users can view a summary table of all available SARS-CoV-2 assays (**Figure 4c**) and download complete datasets for each screen with a single click for further analysis, including compound metadata and MOAs (**Supplemental Table 2**)^21^. A data dictionary describing the contents and format of these datasets in greater detail is provided (**Table 4**). By clicking on individual assays, users can access a view containing detailed assay protocols to allow independent evaluation and confirmation of method and compound activities, as well as details on data normalization, assay interpretation and per-plate assay performance metrics by library (**Figure 4d**).

**Table 4.**
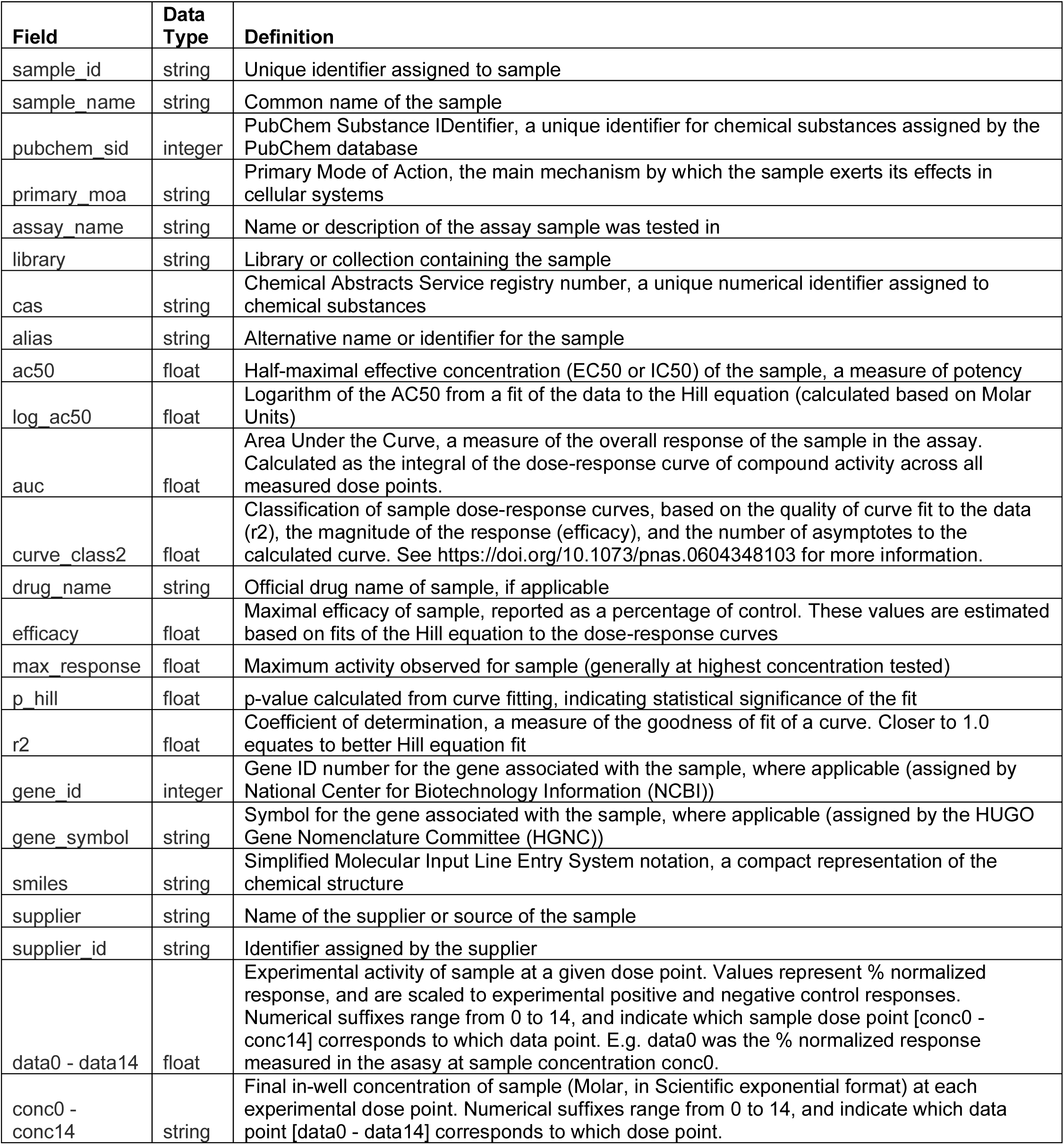
Data dictionary for OpenData Portal screening datasets. Definitions of the columns and data types included in the OpenData Portal screening datasets.

| Field | Data Type | Definition |
| --- | --- | --- |
| sample_id | string | Unique identifier assigned to sample |
| sample_name | string | Common name of the sample |
| pubchem_sid | integer | PubChem Substance IDentifier, a unique identifier for chemical substances assigned by the PubChem database |
| primary_moa | string | Primary Mode of Action, the main mechanism by which the sample exerts its effects in cellular systems |
| assay_name | string | Name or description of the assay sample was tested in |
| library | string | Library or collection containing the sample |
| cas | string | Chemical Abstracts Service registry number, a unique numerical identifier assigned to chemical substances |
| alias | string | Alternative name or identifier for the sample |
| ac50 | float | Half-maximal effective concentration (EC50 or IC50) of the sample, a measure of potency |
| log_ac50 | float | Logarithm of the AC50 from a fit of the data to the Hill equation (calculated based on Molar Units) |
| auc | float | Area Under the Curve, a measure of the overall response of the sample in the assay. Calculated as the integral of the dose-response curve of compound activity across all measured dose points. |
| curve_class2 | float | Classification of sample dose-response curves, based on the quality of curve fit to the data (r2), the magnitude of the response (efficacy), and the number of asymptotes to the calculated curve. See https://doi.org/10.1073/pnas.0604348103 for more information. |
| drug_name | string | Official drug name of sample, if applicable |
| efficacy | float | Maximal efficacy of sample, reported as a percentage of control. These values are estimated based on fits of the Hill equation to the dose-response curves |
| max_response | float | Maximum activity observed for sample (generally at highest concentration tested) |
| p_hill | float | p-value calculated from curve fitting, indicating statistical significance of the fit |
| r2 | float | Coefficient of determination, a measure of the goodness of fit of a curve. Closer to 1.0 equates to better Hill equation fit |
| gene_id | integer | Gene ID number for the gene associated with the sample, where applicable (assigned by National Center for Biotechnology Information (NCBI)) |
| gene_symbol | string | Symbol for the gene associated with the sample, where applicable (assigned by the HUGO Gene Nomenclature Committee (HGNC)) |
| smiles | string | Simplified Molecular Input Line Entry System notation, a compact representation of the chemical structure |
| supplier | string | Name of the supplier or source of the sample |
| supplier_id | string | Identifier assigned by the supplier |
| data0 - data14 | float | Experimental activity of sample at a given dose point. Values represent % normalized response, and are scaled to experimental positive and negative control responses. Numerical suffixes range from 0 to 14, and indicate which sample dose point [conc0 - conc14] corresponds to which data point. E.g. data0 was the % normalized response measured in the assay at sample concentration conc0. |
| conc0 - conc14 | string | Final in-well concentration of sample (Molar, in Scientific exponential format) at each experimental dose point. Numerical suffixes range from 0 to 14, and indicate which data point [data0 - data14] corresponds to which dose point. |

The web application framework used to upload and disseminate screening datasets on the OpenData Portal website is described in **Figure 5**. The screening data is first generated, normalized, and processed in-house at NCATS. It is then deposited in a PostgreSQL database, which is served via API to the Angular-based web client. Users can explore and download the data through the OpenData Portal web interface, with D3.js-powered visualizations providing interactive charts for enhanced data exploration.

**Figure 5.**
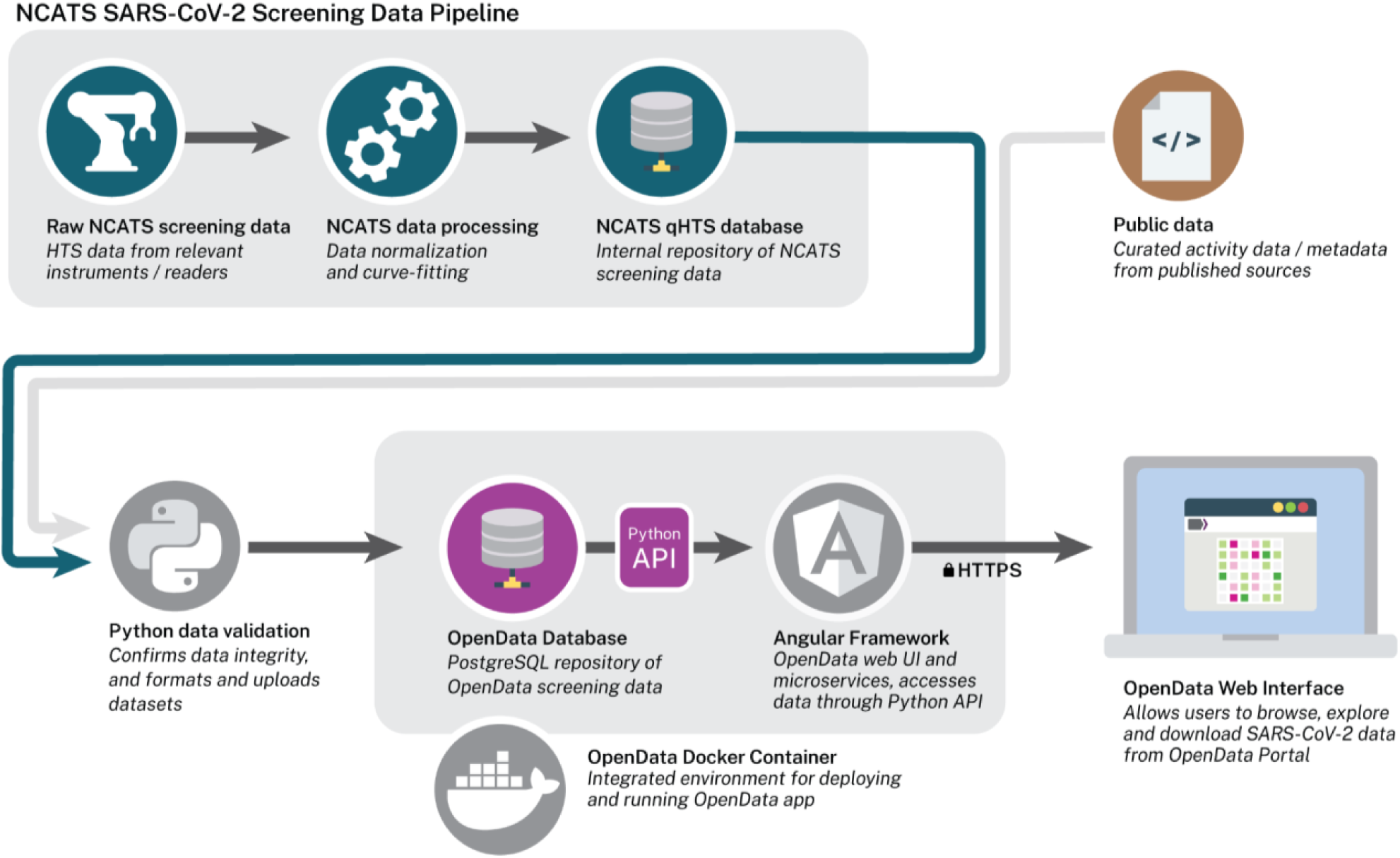
OpenData Portal Web Application Framework. Raw NCATS quantitative high-throughput screening data is normalized using assay-relevant controls. Curve-fitting algorithms are applied, providing curve classes and estimates of efficacy and potency for all compounds. Processed data is written to internal NCATS qHTS databases for access. Screening datasets flagged for public access via the OpenData Portal are first validated using custom Python scripts to further unify formatting of data and metadata, then deposited into a dedicated OpenData Portal Postgres database that can be accessed via API. An Angular-based web client consumes this API to allow exploration of the screening data, which can be previewed and downloaded from the OpenData Portal web interface. D3.js-based data visualizations have also been built, which allow users to explore intuitive and interactive charts designed to surface specific insights of interest to the SARS-CoV-2 community. Subsequent expansion of the OpenData Portal has enabled the sharing of public data (e.g. reported therapeutic activity against SARS-CoV-2 variants) curated by NCATS scientists; these datasets are similarly validated via Python script, deposited into the Postgres database, and shared on the OpenData Portal for visualization, exploration and download.

## Discussion

The NCATS OpenData Portal is premised on the idea that rapid and open sharing of complete datasets will accelerate the process of research and discovery, consistent with NCATS’s mission to advance translational science. The experience of sharing SARS-CoV-2 screening datasets – including inactive compounds and matched counterscreen data – publicly, and in close to real-time, has borne out this idea.

During the pandemic, the scientific community generated an abundance of therapeutic hypotheses against COVID-19, many centered around potential drug repurposing opportunities, natural product-based therapies, or other mechanistic classes of interest. Preliminary SARS-CoV-2 screening efforts conducted at NCATS and shared through the OpenData Portal canvassed much of this chemical and biological space, with over 10,000 approved drugs and well-characterized compounds (**Table 2**) tested against more than a dozen SARS-CoV-2 targets and assays, spanning both viral and host targets (**Table 1**, **Figure 1**). The resulting data was shared rapidly and freely with the global community, representing some of the earliest and most comprehensive screening datasets available during the COVID-19 pandemic. NCATS directed scientific queries to the OpenData Portal data browser to provide immediate evidence for the evaluation of early hypotheses, with a view to reducing duplicative screening work, focusing development efforts on the most promising candidates, and providing a forum for the verification of scientific observations.

The OpenData Portal website was designed to encourage exploration of these SARS-CoV-2 datasets and make the results accessible to a general science audience, while enabling data reuse by those in the data science community for developing and validating theories and models (**Figure 4**). This large dataset has been reused a number of times by groups outside of NCATS, not only to generate or refine SARS-CoV-2-related therapeutic hypotheses, but also to train and develop predictive models for potential SARS-CoV-2 therapeutics, and to experimentally validate predicted hits from virtual screens, protein docking, and transcriptome models^22–41^.

### SARS-CoV-2 Screening Results

The SARS-CoV-2 screening assays shared through OpenData Portal covered a wide panel of COVID-19-relevant drug space (**Table 2**). Most of the assays shared in the OpenData Portal (11/17) were cell-based (**Figure 1b-d**), which generally reflects the use of host cell lines as a proxy in viral infection assays. Of these cell-based screens, over half (6/11) measured compound toxicity in host cells as a counterscreen to identify potential off-target toxicity or broader effects on cell growth which could confound interpretation of activity in the primary screens. The live SARS-CoV-2 CPE assay and viral entry assays (those interrogating the spike-ACE2 binding interaction) were among the earliest assays screened and shared, due in part to 1) the early availability of BSL3 CPE models through research contracts, and 2) the scientific community’s strong focus on SARS-CoV-2 spike protein as a drug target early in the pandemic. Complex assays (such as the SARS-CoV-2 pseudotyped particle assay) and biochemical assays utilizing recombinant protein (such as the RdRp polymerase and 3CL and TMPRSS2 protease assays) were generally developed later as custom expression systems and/or limiting research reagents became available.

Although a deeper analysis of the screening hits obtained from this panel of SARS-CoV-2 assays is outside the intended scope of this paper, some broader trends can be observed. In general, the hit rates of high confidence actives observed in the primary screens fell within historically observed qHTS ranges (**Figures 2a-b**)^42^. Assays with higher hit rates tended to have concomitantly high hit rates in corresponding counterscreen assays (e.g. the spike-ACE2 AlphaLISA and SARS-CoV-2 pseudotyped particle assays), suggesting many hits were assay artifacts or exhibited polypharmacology (**Figure 2c**). Such cases further highlight the importance of sharing counterscreen assay datasets for proper interpretation by the scientific community, so that potential false positive activity can be considered. This is especially important when using activity data for model building or validation.

While replication studies are often the next step after a screening campaign, several of the screens shared on OpenData Portal included natural replication experiments (**Table 3**). In general, confirmation rates were inversely correlated to overall screening hit rates; screens with lower hit rates yielded higher replicate confirmation rates, and vice versa. This is likely, in part, a reflection of assay and/or biological stringency, and partly related to the fact that screens with a higher proportion of inactives will have a lower bar for demonstrating confirmed activity across replicates: in quantitative high-throughput screening, high or low confidence actives must reproduce both curve goodness-of-fit and efficacy (which are used to define curve classes), whereas inactive compounds need only fail to reproduce one or the other of those above standards to qualify as inactives. In any case, the high observed replicate confirmation rates provide further evidence of the robustness of these assays and increase confidence in the quality of the activity datasets shared on the OpenData Portal.

When comparing high confidence actives across screens (**Figure 2d**), there appeared to be the greatest overlap between the pseudotyped particle assays (SARS-CoV-2 and MERS in particular), which share similar assay protocols. Intriguingly, the spike-ACE2 AlphaLISA assay shared a number of hits in common with the pseudotyped particle assays, suggesting there may be some broader effects on viral entry, despite the fact that MERS does not engage human ACE2 for host cell entry^43^. In general, most high confidence actives were quite assay-specific, with observed activity only in a single screen.

### Importance of Complete Data-sharing

Due to the scale and urgency of early efforts against COVID-19, traditional channels for disseminating research findings quickly became insufficient to deal with the pace of SARS-CoV-2 research. Though the field experienced an encouragingly rapid and widespread adoption of preprint publications and other open science approaches, a data transparency gap has persisted in many screening publications^44^. Hit-to-lead discovery publications often only report active compounds or a cherrypicked subset of hits without disclosing complete screening datasets, particularly in the private sector where the intellectual property value of proprietary screening collections can limit structure disclosure and data sharing^45,46^. Inactive or weakly active compounds can comprise >99% of compounds tested in many screens, and active compounds with identified counterscreen activity are often disclosed without this additional data and context or are excluded from publication altogether^42,47^. As such, the research community is unable to access the majority of experimental screening results.

The NCATS OpenData Portal was designed to address these gaps in transparency and speed, and in practice, the site successfully demonstrated that this open approach can be implemented at scale. Nearly 90% of the complete SARS-CoV-2 screening datasets generated at NCATS were shared on the site before or within 2 weeks of preprint or peer-reviewed publication release, and most (59%) were shared in advance of publication (**Figure 3a**). The majority of this data (13/17 screens) was released within 2020, the first year of the COVID-19 pandemic, and all released datasets included concentration-response data on screened compounds, including all inactive compounds (**Figure 3b**).

Given the widespread interest in drug repurposing, the slow, incomplete or inaccessible disclosure of approved drug screening data leads to unnecessary duplication of efforts throughout the global research community. The ultimate cost of duplicative and poorly-designed COVID-19 clinical trials has been well documented, and more broadly, an estimated $10 to $50 billion is spent each year on studies that cannot be successfully reproduced^48,49^. Many large and expensive research campaigns have undoubtedly been conducted based on incorrect therapeutic hypotheses, and it is likely that many of these hypotheses could have been rejected earlier if SARS-CoV-2 screening data - specifically negative data - had been shared more widely or efficiently.

Furthermore, while drug repurposing screening campaigns are popular starting points for therapeutic development, repurposing success stories are notoriously rare^50^. This is not to say that repurposing campaigns are not worth the effort; indeed the cost of a repurposing screen could be considered trivial compared to the value of a significantly shortened path from discovery to drug approval, and significant biological insight can be derived based on pharmacogenomic data^51^. However, given the low success rate of repurposing screening, it follows that most of the value of these campaigns lies in what the broader scientific community can learn from their complete datasets. Though negative data may be uninteresting from the perspective of a single screening campaign, it can be critical for understanding and validating disease and drug mechanisms-of-action. For example, a drug with a known mechanism may show early activity and lead to the conclusion that modulation of that gene or drug target is critical for a phenotype; however, a full screening dataset may demonstrate that other drugs with the same mechanism are inactive. In this respect, negative data is critically important to support (or temper) hypothesis generation.

To this end, artificial intelligence and machine learning (AI/ML) modeling holds significant promise to accelerate discovery in drug development. However, these techniques are entirely dependent on the availability of well-annotated, high-quality activity data to train and refine AI/ML models^52^. This is particularly true with respect to negative data, which is shared less consistently than positive activity data, but which is equally critical to improve the accuracy of such predictive bioactivity models. The more than 400,000 data points of SARS-CoV-2 screening data shared on the NCATS OpenData Portal were used by a number of research groups to rapidly build and test predictive models of SARS-CoV-2 therapeutics and drug repurposing opportunities, underscoring the value of this comprehensive data-sharing approach^24,25,28–30,32–37,39–41^.

### Future directions

Soon after the initial SARS-CoV-2 screening datasets were shared on the OpenData Portal, in coordination with the NIH’s Accelerating COVID-19 Therapeutic Interventions and Vaccines (ACTIV) public-private partnership, the site was significantly expanded to curate and disseminate *in vitro* therapeutic data reported against known and emerging SARS-CoV-2 variants during the pandemic^53^. The Portal maintained one of the most comprehensive public databases of *in vitro* therapeutic activity data available against known variants to help guide decision-making in government and industry^54^. The Portal continued to adapt as the therapeutic landscape of COVID-19 shifted, and additional databases and pages were launched to track different therapeutic development data, including relative activity of COVID-19 booster regimens, development status of potential COVID-19 therapeutics, summaries of animal models and published *in vivo* challenge studies modeling SARS-CoV-2 infection, and summaries of real-world evidence studies on EUA/FDA approved and revoked COVID-19 therapeutics^55–60^.

The OpenData Portal was built with a flexible framework (**Figure 5**) to allow the site to expand and share additional data types and structures, and to act as a disease agnostic data-sharing hub for NCATS beyond COVID-19. Future updates are focused on improving the accessibility and openness of OpenData Portal datasets to further enable data reuse and ease interpretation. These efforts include the implementation of documented APIs to enable users to programmatically retrieve OpenData Portal datasets and allow integrated data connections with other sites and tools. This future work is designed to allow the OpenData Portal resource to efficiently scale as existing data sources grow, and to expand in scope as new disease types and data sources are incorporated.

In January 2023, the National Institutes of Health (NIH) issued a mandate requiring most of the institutions it funds annually - as well as its own intramural research laboratories - to include a data management plan in their applications and annual reports, and to eventually make their data publicly available^61^. While the OpenData Portal was launched in advance of this mandate, this resource aims to set a standard for open science and data-sharing in the government and significantly shorten the timeline on which NCATS screening data is shared. In summary, the OpenData Portal described herein was designed to share NCATS SARS-CoV-2 complete datasets openly, without restriction, and in real-time given the urgency of the COVID-19 pandemic response. Going forward, it provides a scalable, established platform to enable rapid sharing of NCATS’s own early discovery screens, experimental datasets, protocols, predictive models and beyond - regardless of disease-focus – with the goal of accelerating cures to patients.

## Methods

### SARS-CoV-2 Screening Assays

The high-throughput SARS-CoV-2 screening assays and counterscreens shared on the OpenData Portal were developed at NCATS. Protocols and screening performance metrics for individual assays are available on the website from the SARS-CoV-2 Assays Page (https://opendata.ncats.nih.gov/covid19/assays). Detailed protocols, descriptions of the development of these assays, and expanded discussions of individual screen results can be found in assay-specific publications on those efforts^62–69^ (**Table 1**).

### Compound Libraries

The NCATS Pharmaceutical Collection (NPC) was assembled at NCATS and contains 2,678 drugs approved by the US FDA and foreign health agencies in the European Union, United Kingdom, Japan, Canada, and Australia, as well as some compounds tested in human clinical trials^15,16^. The NCATS Pharmacologically Active Chemical Toolbox (NPACT) is a library of annotated compounds that inform on novel phenotypes, biological pathways and cellular processes. Compounds in the NPACT library represent more than 7,000 mechanisms and phenotypes identified in the literature and worldwide patents, and cover biological interactions within mammalian, microbial, plant and other model systems. The NCATS Mechanism Interrogation Plate (MIPe) library contains 2,477 mechanism-based bioactive compounds, targeted against more than 860 mechanisms of action^70,71^. The Anti-infective Library is a set of compounds curated by NCATS scientists early in the pandemic to test potential repurposing candidates against coronavirus infection. It contains compounds from the scientific literature with reported anti-infective activity, including antibacterials, antivirals, antifungals, and antiparasitics. In general, library compounds were dissolved as 10 mM DMSO stock solutions and titrated at 1:5 dilution for primary screens at 4 or more concentration points. All libraries make use of interplate titrations - where compound titrations are spread across multiple assay plates - which minimizes the effect of any single plate performance issues, as activity curves can still be generated from the remaining plates/titration points.

### Data Analysis

Analysis methods for each screening assay are described in detail in assay-specific publications. In general, raw concentration–response data for each screened compound was fitted using a four-parameter logistic model, yielding potency (IC_50_), efficacy (maximal response) and r^2^ goodness-of-fit values, which were further used to bin compounds into categorical curve classes. For screens where only four concentrations were tested, curves were fitted using a three-parameter logistic model. Raw plate reads for each concentration were normalized relative to neutral controls (DMSO) and assay-relevant positive controls (equivalent to 0% activity, or complete inhibition) and negative controls (equivalent to 100% activity, or basal/uninhibited activity), and normalized data was subsequently corrected for plate effects using pattern correction procedures as appropriate^72^. Assay control well activity was assessed, and Z-scores and signal-to-background (S:B) metrics were calculated for each plate and manually reviewed to ensure assay metrics fell within an expected range, given each assay’s historical performance.

For the purposes of this study, active compounds were defined as either high confidence or low confidence hits. High confidence actives were compounds with either 1) dual asymptotic curve fits with r^2^ ≥ 0.9 or 2) single asymptotic curve fits with r^2^ > 0.9 and efficacy > 80%, which fall into the 1.1, 1.2, 2.1 and 5 (unclassified activity) curve classes. Low confidence actives were compounds with 1) dual asymptotic curve fits with r^2^ < 0.9 and efficacy ≥ 80% or 2) single asymptotic curve fits with at least single point activity, classified as 1.3, 1.4, 2.2, 2.3, 2.4 or 3 (demonstrating inhibition, rather than activation) curve classes (https://opendata.ncats.nih.gov/covid19/curveclasses)^19^. For the CPE assay (assay #1), which measures the rescue of virus-induced cell death as an increase in cell viability, compounds which demonstrated positive curves (activation-like) were considered active, and compounds with negative (inhibition-like) or class 4 (no activity) curves were considered inactive. For all other assays (#2-17), compounds with negative curves were considered active, while compounds with positive or class 4 curves were considered inactive.

### OpenData Portal Website and Interface

The OpenData Portal, which can be accessed at https://opendata.ncats.nih.gov/, is intended to provide clinicians and researchers with a user-friendly tool that allows direct comparison of compounds across multiple SARS-CoV-2 assays. Quantitative high-throughput screening (qHTS) data and detailed protocol information are available for every assay screened, and all primary concentration-response data are made freely available through direct download. The OpenData Portal web application was built using Angular v.16 (https://angular.io), a TypeScript-based single-page web application framework (**Figure 5**). The client-side application utilizes a Python v.3 (https://www.python.org/) high-performance API to pull validated data from the OpenData Portal PostgreSQL database and enable interactive operations on the underlying datasets. Complex interactive charts were created on the site using D3.js v.7 (https://d3js.org) and were designed to surface specific insights of interest to the SARS-CoV-2 community. Users can explore and filter these complex datasets to find compounds, assays and activities of interest.

## Data availability

NCATS SARS-CoV-2 normalized screening data and compound metadata are dedicated to the public domain under the Creative Commons CC0 1.0 Universal Public Domain Dedication, and are available for download on the NCATS OpenData Portal site at https://opendata.ncats.nih.gov/covid19/assays. Users can opt to download the full SARS-CoV-2 screening dataset (all 17 assays) or individual assay datasets. Documentation of the assay methods are available on the site, and a data dictionary with column definitions is packaged with data downloads.

In addition to the data portal site, all source data supporting the findings of this study are available within the paper and its Supplementary Information.

## Code availability

No code is shared in this paper.

## Author Contributions

K.R.B., M.D.H., T.Z., M.S., and D.C.L. contributed to the conceptualization and design of the resource; B.B., C.Z.C., K.G., Q.M.H., S.C.K., E.M.L., M.P., A.R., J.H.S., M.X., W. Zhu, M.X., Y.F., M.F., W. Zheng, A.S. and M.D.H. designed, conducted and/or directed the SARS-CoV-2 assays, high-throughput screens and supporting experiments to yield the original shared datasets; C.K.T., Z.I., S.M., P.S. and K.M.W. provided experimental support and access to critical screening/experimental core capabilities; L.C., H.G., T.E., X.H., A.V.Z., A.P. and E.A.M. contributed data analysis and other informatics-related support; T.Z., K.D., A.F., B.N., N.H.,T.M., K.W., M.B., S.M. and A.V provided web/software development and IT operations supports; K.R.B., P.C.S, Y.C., D.P.D., K.D., B.J.P., T.N.T. and K.K.W. provided project coordination and program support; K.R.B., P.S.C., and R.T.E. prepared the manuscript with the assistance and feedback from all other co-authors.

## Acknowledgements

We thank the NCATS DPI scientists for bravely sharing data more quickly and openly than ever in the interest of helping the research community tackle the COVID-19 pandemic. We also thank the NCATS laboratory support staff, Compound Management and Automation groups, Informatics and IT technical teams for helping enable this screening work on short timelines and under challenging lockdown conditions. We additionally thank our colleagues in the ACTIV public-private partnership and the ACTIV TRACE working group for their invaluable feedback on the OpenData Portal website, and their help raising awareness about the resource.

## Supplemental Tables

**Supplemental Table 1. Anti-infective library composition.** Summary of the names and associated metadata of the small molecules in the NCATS Anti-infective Library assembled and screened for the OpenData Portal.

**Supplemental Table 2. Complete SARS-CoV-2 screening dataset.** Normalized data and metadata collected for all SARS-CoV-2 screening datasets shared on the OpenData Portal.

